# Emerging Antifungal Resistance in Falco Species: A Novel Model for Human Medicine

**DOI:** 10.1101/2023.03.22.533837

**Authors:** Sibi Das, Sethi Das Christu, Christudas Silvanose, Ambili Binoy, Panagiotis Azmanis, Antonio Di Somma, Jibin V Gladston

**Author notes:** For correspondence: Dr. Sibi Das, Sri Siddhartha Medical College, Tumkuru, Karnataka, India.

## Abstract

Antifungal resistance is a growing concern in the medical community, as fungal infections are becoming increasingly difficult to treat. In this study, Falco species were used as novel models for studying antifungal resistance since Aspergillosis, a fungal disease is common in falcons. The most isolated fungi in this study were *A. fumigatus, A. flavus, A. niger*, and *A. terreus*, all of which can cause aspergillosis in falcons. Isavuconazole, posaconazole, and voriconazole had the lowest MICs among the drugs tested, suggesting that they may be effective treatment options. However, this study showed that 34% of the isolates were resistant to itraconazole, which is an increase from 21% in 2006. There is no resistance to voriconazole found in 2006 and 2011, but a 9% resistance rate was noted in 2022. Similarly, there is no resistance to posaconazole and isavuconazole was noticed in 2011, but resistance of 4.7% and 5.8%, respectively was noticed in 2022. Amphotericin B, which showed a 51% resistance rate in 2006, became even more resistant with an 80% rate in 2011, leading to its discontinuation from the treatment of falcons against aspergillosis. This study highlights a significant rise in antifungal resistance, which is a challenging problem in both falcon and human medicine.

## Introduction

The emergence of resistant fungal pathogens posed a significant threat to antifungal treatments, particularly for immunocompromised patients and individuals with chronic conditions. To better understand antifungal resistance during treatment and develop new strategies for combating fungal infections, there was increasing interest in studying animal models. Falcons (Falco species) were particularly susceptible to fungal infections, including aspergillosis, which was a common cause of morbidity and mortality among falcons [1-4]. Recent studies had reported the emergence of antifungal resistance in falcons, making them an interesting model for studying antifungal resistance mechanisms [5].

Antifungal resistance in falcons was an increasingly concerning issue in avian medicine. Aspergillosis was caused by Aspergillus species, a fungal pathogen that could colonize the respiratory system and cause respiratory distress, chronic infections, and even death in severe cases [6]. Overuse or inappropriate use of antifungal drugs in captive birds, as well as the spread of resistant fungal strains between birds in falconry or rehabilitation centers, could lead to antifungal resistance in falcons [7]. The study of antifungal resistance in falcons was essential for several reasons. Firstly, falcons were a valuable animal model for studying fungal infections and host-pathogen interactions. Secondly, the emergence of antifungal resistance in falcons could have implications for managing fungal infections in other animal species, including humans. Lastly, studying antifungal resistance in falcons could offer insights into resistance during treatment, which could identify new therapeutic targets for both animal and human medicine. Fungal diseases had emerged in association with post-COVID-19, as the COVID-19 virus weakened the immune system, making individuals more susceptible to fungal infections. Thus, it was crucial to develop new and effective antifungal treatments to address this growing concern.

This manuscript investigated the emerging antifungal resistance in falcons, with a particular focus on triazole resistance, and its potential implications for human medicine. Thus, compared the findings with existing data on human antifungal resistance to identify potential areas of overlap and divergence. By studying the incidence and prevalence of aspergillosis in falcons, which was higher than any other species, advanced diagnosis and treatment procedures had been developed in Falcon medicine, including in-vivo and in-vitro MIC antifungal studies. The project had the potential to provide important insights into the mechanisms of antifungal resistance and to identify novel therapeutic targets that could be applied to both animal and human medicine. By examining a non-traditional model such as falcons, we could also gain a new perspective on the evolution and spread of antifungal resistance and contribute to the development of innovative and effective strategies for managing fungal infections in both animals and humans.

## Materials and methods

Biopsy samples were collected during the endoscopy of air sacs of eighty-six falcons which include Peregrine falcons (*Falco peregrinus*), Saker falcons (*Falco cherrug*), Gyr (*Falco rusticolus*), and hybrid falcons such as Gyr x Peregrine (*F. rusticolus x F. peregrinus*) and Gyr x Saker (*F. rusticolus x F. cherrug*), affected with lower respiratory tract fungal infection. Samples were cultured in Sabouraud’s chloramphenicol agar (SCA) and incubated at 37°C for 3-5 days. Fungi were identified by culture appearance and morphological characteristics under the microscope using lactophenol aniline blue stain preparation. Antifungal studies were done in RPMI media using antifungal MIC E-test strips (bioMerieux, France) and the plates were incubated at 37° C in an incubator for 48 hours. The Minimum Inhibitory Concentration (MIC) was recorded as the lowest concentration of the antifungal agent that inhibited fungal growth. The first phase of the study was done in 2006 with 100 isolates of *Aspergillus* species, the second phase of the study was done in 2011 with 117 isolates of *Aspergillus* species, and the current study, the third phase was done in 2023 with 86 fungal isolates to compare the evolution of antifungal resistance in the past 15 years.

## Results

Aspergillosis was confirmed through endoscopic examinations, cytological examinations, and mycology cultures. During an endoscopy, characteristic signs of aspergillosis may include the presence of nodules or growths on the lining of the respiratory tract (Figure 1). These growths can appear as whitish-yellow patches or raised bumps (Figure 2).

**Figure 1.**
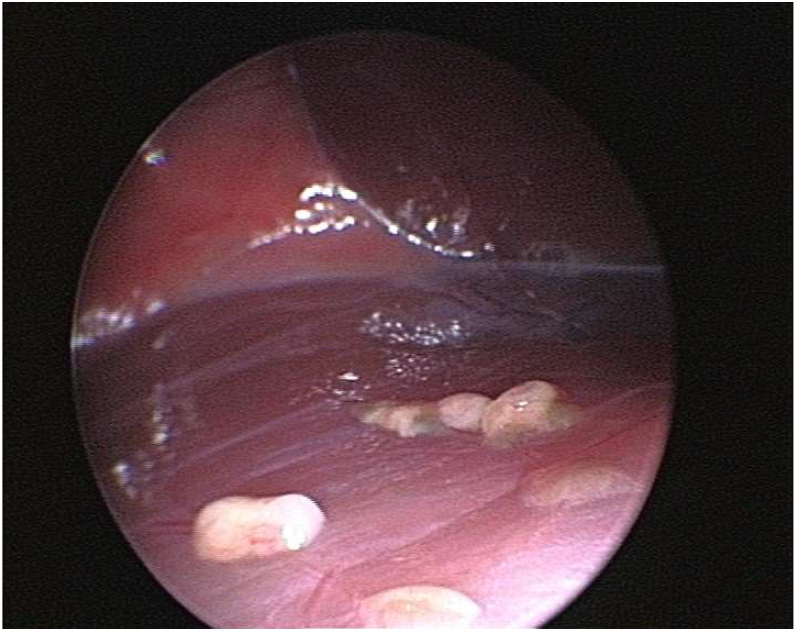
Endoscopic view of multiple aspergillomas showing multiple nodular raised growth in the air sac of a Gyrfalcon.

**Figure 2.**
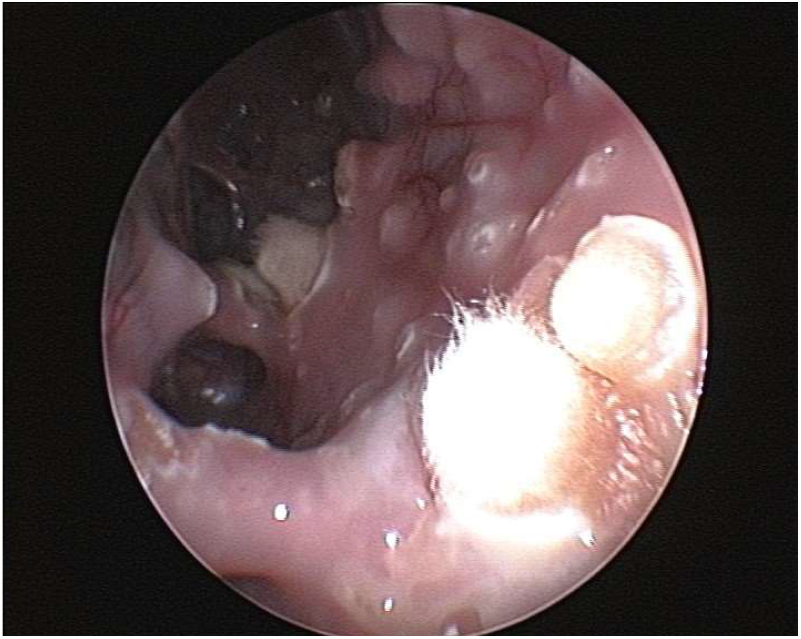
Endoscopic view of multiple aspergillomas with few sporulating aspergillomas and raised bumps in the air sac of a Gyrfalcon.

The diagnosis of aspergillosis through cytology includes the appearance of fungal elements and cellular changes depending on the site of infection (Figure 3). Confirmation of aspergillosis was through fungus culture (Figure 4) and microscopic examination of the culture under lactophenol blue stain preparation for the morphology of conidiophores and hyphae.

**Figure 3.**
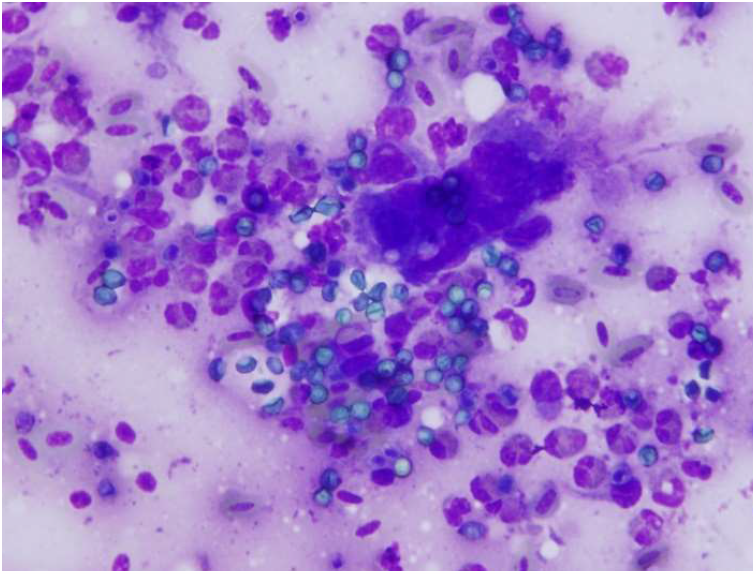
Cytology imprint smear of the biopsy sample collected from the air sac of a falcon with aspergillosis. Note the giant cell formation and fungal spores in an inflammatory cell background.

**Figure 4.**
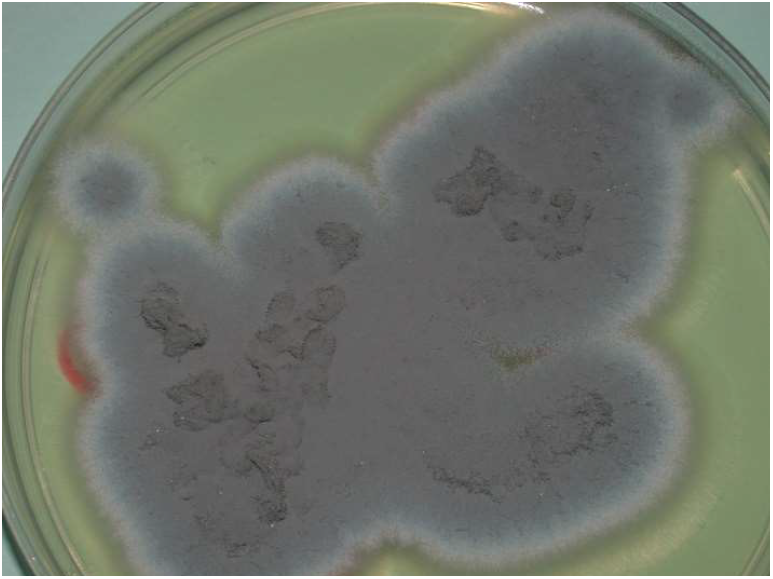
Culture appearance of *Aspergillus fumigatus* isolated from the air sac of Gyr falcon with aspergillosis showing bluish colonies after 72 hours of incubation.

The minimum inhibitory concentration (MIC) value represents the lowest concentration of a drug required to inhibit the isolate, and the lowest MIC value indicates greater effectiveness against the fungal isolate. Table 1 reports the median MIC values and range of MIC values for each antifungal agent tested against each fungal species obtained from the air sac of falcons.

**Table 1.** MIC of fungal isolates from the air sac of falcons with ≤ 1μg/ml

| Fungi isolated (n)* | POZ | ISZ | VOZ | ITZ |
| --- | --- | --- | --- | --- |
|  | Median (Range) | Median (Range) | Median (Range) | Median (Range) |
| **A. fumigatus (36)** | 0.3 (0.094-1) | 0.19 (0.032-1) | 0.19 (0.032-1) | 1 (0.25-1) |
| **A. flavus (13)** | 0.75 (0.125-1) | 0.38 (0.25-1) | 0.025 (0.032-0.75) | 0.75 (0.38-1) |
| **A. niger (18)** | 0.38 (0.125-1) | 0.25 (0.125-0.75) | 0.25 (0.125-1) | 0.75 (0.2-1) |
| **A. terreus (14)** | 0.38 (0.19-0.75) | 0.38 (0.064-0.75) | 0.5 (0.094-0.75) | 0.75 (0.25-1) |
| **A. nidulans (2)** | 0.19 (0.19-1) | 0.064 (0.02-0.125) | 0.094 (0.047-0.125) | 0.5 (0.38-1) |
| **A. versicolor (1)** | 0.25 (0.25) | 0.023 (0.023) | 0.064 (0.064) | 0.25 (0.25) |
| **Mucor species (2)** | 0.75 (0.75) | 0.875 (0.25-1) | 0 | 0.625 (0.25-1) |

\*n= total number of isolates, POZ = Posaconazole, ISZ = isavuconazole, VOZ = voriconazole, ITZ = Itraconazole

*A. fumigatus* was the most frequently isolated fungal species, with a total of thirty-six isolates. *A. niger* and *A. terreus* were isolated with a total of 18 and 14, respectively. Posaconazole was effective against *A. fumigatus*, with a median MIC value of 0.3 μg/ml, while isavuconazole and voriconazole showed median MIC values of 0.19 μg/ml against *A. fumigatus*. Itraconazole was less effective, with a median MIC value of 1 μg/ml, and values above 1 μg/ml are considered resistant.

Figure 6 shows the e-test MIC readings of *A. fumigatus* against voriconazole and itraconazole as a typical example. *A. flavus* was isolated from 13 cases, and Posaconazole and itraconazole showed similar activity against *A. flavus*, with median MIC values of 0.75 μg/ml and 0.75 μg/ml, respectively. Isavuconazole and voriconazole were more potent, with median MIC values of 0.38 μg/ml and 0.025 μg/ml, respectively, against *A. flavus*.

**Figure 6.**
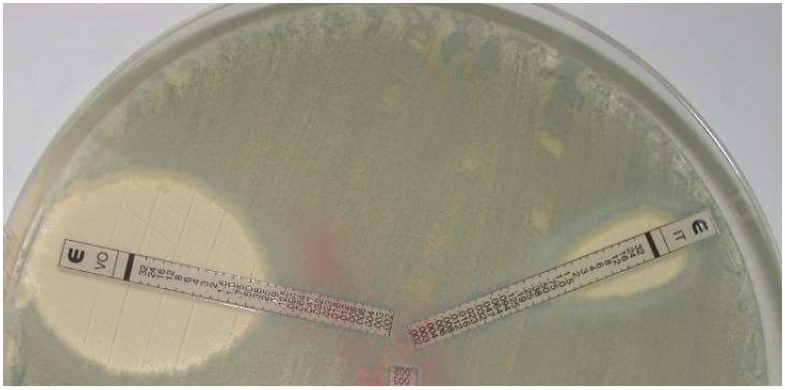
E-test reading of *A. fumigatus*, an isolate from the air sac of a Gyrfalcon showing voriconazole with 0.38 μg/ml and itraconazole with 1 μg/ml.

The results of the study showed that posaconazole and isavuconazole were effective against *A. niger*, with median MIC values of 0.38 μg/ml and 0.25 μg/ml, respectively. Voriconazole also demonstrated good activity against *A. niger*, with a median MIC value of 0.25 μg/ml, while itraconazole showed a higher median MIC value of 0.75 μg/ml. A. terreus was most susceptible to isavuconazole and posaconazole, with a median MIC value of 0.38 μg/ml. Voriconazole showed similar activity against *A. terreus*, with a median MIC value of 0.5 μg/ml, while itraconazole had a higher median MIC value of 0.75 μg/ml. *A. nidulans* and A. versicolor showed good antifungal activity against all antifungals, but they were not commonly isolated. The two isolates of *Mucor* species were sensitive to posaconazole and itraconazole, with median MIC values of 0.75 μg/ml and 0.625 μg/ml, respectively. Isavuconazole showed a higher MIC value of 0.875 μg/ml, while voriconazole was found to be resistant against *Mucor* species.

Table 2 shows the mic of antifungal drugs against fungal isolates from the air sac of falcons with >1 μg/ml and it falls under the resistant level in birds. A. fumigatus isolated from 3 cases was resistant to posaconazole and isavuconazole with a median MIC of 1.5 μg/ml, 4 cases were resistant to voriconazole with a median MIC of 3 μg/ml, and 15 isolates were resistant to itraconazole with a median MIC of 3 μg/ml. The resistance of A. flavus includes one isolate to voriconazole and 3 isolates to itraconazole with MIC of 1.5 μg/ml. Each isolate of *A. niger* was resistant to isavuconazole and voriconazole with a MIC of 1.5 μg/ml, while 8 isolates were resistant to itraconazole with a median MIC of 4 μg/ml. *A. terreus* resistance was noticed in one case to posaconazole with 1.5 μg/ml, isavuconazole and voriconazole with 32 μg/ml; while 3 isolates of *A. terreus* were resistant to itraconazole with a median MIC was 1.5 μg/ml. Mucor species were isolated from two cases that were resistant to voriconazole with a median MIC > 32 μg/ml.

**Table 2.** MIC of fungal isolates from the air sac of falcons with > 1μg/ml

| Fungi isolated | POZ | ISZ | VOZ | ITZ |
| --- | --- | --- | --- | --- |
|  | Median (Range) | Median (Range) | Median (Range) | Median (Range) |
| ***A. fumigatus*** | 1.5<br>(1.5 -4)<br>n = 3 | 1.5<br>(1.5-32)<br>n = 3 | 3<br>(1.5 -32)<br>n = 4 | 3<br>(1.5 -32)<br>n = 15 |
| ***A. flavus*** | n = 0 | n = 0 | 1.5<br>(1.5)<br>n = 1 | 1.5<br>(1.5 -2)<br>n = 3 |
| ***A. niger*** | n = 0 | 1.5<br>(1.5)<br>n = 1 | 1.5<br>(1.5)<br>n = 1 | 4<br>(2-32)<br>n = 8 |
| ***A. terreus*** | 1.5<br>(1.5)<br>n = 1 | 32<br>(32)<br>n = 1 | 32<br>(32)<br>n = 1 | 1.5<br>(1.5 -2)<br>n = 3 |
| ***A. nidulans*** | n = 0 | n = 0 | n = 0 | n = 0 |
| ***A. versicolor*** | n = 0 | n = 0 | n = 0 | n = 0 |
| ***Mucor* species** | n = 0 | n = 0 | 32<br>(32)<br>n = 2 | n = 0 |
| **Total** | n = 4 | n = 5 | n = 8 | n = 29 |

\*n= total number of isolates, POZ = Posaconazole, ISZ = isavuconazole, VOZ = voriconazole, ITZ= Itraconazole

Table 3 shows a comparison of antifungal resistance of Aspergillus species isolated in 2006, 2011, and 2022. These results may have important implications for the treatment and management of fungal infections in birds of prey, as well as for understanding the impact of antifungal use on the emergence of resistance in wildlife and human pathogens.

**Table 3.** Comparison of Antifungal resistance of Aspergillus species isolated from the air sac of falcons in 2006, 2011, and 2022.

| Anti-fungal | 2006 (A)* | 2006 (B)* | 2011 | 2022 |
| --- | --- | --- | --- | --- |
|  | n (Res) | n (Res) | n (Res) | n (Res) |
| **POZ** | - | - | 117 (00%) | 86 (4.7%) |
| **ISZ** | - | - | - | 86 (5.8%) |
| **VOZ** | 45 (00%) | 16 (00%) | 117 (00%) | 86 (09%) |
| **ITZ** | 45 (21%) | 05 (20%) | 117 (06%) | 86 (34%) |
| **AMB** | 45 (51%) | 06 (100%) | 117 (80%) | - |

\*A-Before treatment, B-After treatment, n = number of isolates, AMB = Amphotericin B, POZ = Posaconazole, ISZ = isavuconazole, VOZ = voriconazole, ITZ= Itraconazole

## Discussion

MIC testing is a process of determining the lowest concentration of an antimicrobial drug that can inhibit the growth of a specific microorganism. However, the isolates in this study showed MICs higher than 1 μg/ml, indicating resistance to antifungal drugs. This highlights the need to monitor antifungal resistance in birds and develop new treatment strategies to address this issue.

This study aimed to assess the MICs of various antifungal drugs against fungal isolates obtained from the air sacs of falcons. The first phase of the study was conducted in 2006, the second phase in 2011, and the current study is the third phase, which looks back to understanding the evolution of antifungal resistance over the past 15 years.

The phase 1 study in 2006 reported the MICs of fungi isolated from the air sacs of falcons before and after antifungal treatment. Before treatment, 95% of the isolates, including *A. fumigatus, A. flavus, A. niger*, and *A. terreus*, were susceptible to voriconazole at MICs up to 0.38 μg/ml, and all the isolates were susceptible at MICs up to 1 μg/ml. Before treatment, 21% of the isolates, including *A. fumigatus* (27.6%), *A. flavus* (16.6%), *A. niger* (100%), and *A. terreus* (23%), were resistant (MIC ≥1 μg/ml) to itraconazole. Furthermore, 51% of the isolates, including *A. fumigatus* (31%), *A. flavus* (78%), *A. niger* (14%), and *A. terreus* (77%), had MICs of over 1 μg/mL to amphotericin B. After treatment, their MICs increased significantly (8).

However, the study found no significant differences between the MICs of voriconazole and itraconazole for the different *Aspergillus* species before and after treatment with these antifungal agents. The findings suggest that voriconazole is highly effective against *Aspergillus* species in falcons, with 100% of the isolates being susceptible, while itraconazole was less effective, with 21% of the isolates showing resistance. In contrast, amphotericin B was less effective, with 51% of the isolates being resistant, and after treatment, the resistance of amphotericin B increased significantly (8).

Amphotericin B is an old antifungal drug that has been used in both human and veterinary medicine to treat systemic fungal infections. However, many reports in the literature on falcon medicine have shown that *Aspergillus* species can develop resistance to Amphotericin B, which was a commonly used antifungal drug in the past. This has raised concerns about the drug’s effectiveness in managing fungal infections in falcons, especially Aspergillosis. Additionally, the drug is known to have potential adverse effects, such as renal toxicity, hypokalemia, and infusion-related reactions. In falcons, Amphotericin B use can also result in hepatotoxicity and nephrotoxicity [9, 10]. Furthermore, reports have indicated that Amphotericin B resistance is emerging in falcons, particularly in cases of chronic and recurrent Aspergillosis. Studies have reported the emergence of Amphotericin B-resistant *Aspergillus fumigatus* in captive falcons, leading to treatment failure and poor clinical outcomes [9, 10]. Therefore, Amphotericin B is no longer used in falcon medicine for the treatment of Aspergillosis.

Phase 2 study was conducted in 2011 with 117 Aspergillus species isolates. All isolates were found to be sensitive to voriconazole and posaconazole, while 6% of the isolates were resistant to itraconazole, and 80% of the isolates were resistant to Amphotericin B. Furthermore, 86% of the isolates were resistant to ketoconazole, with a MIC> 1μg/ml. All isolates were resistant to 5-flucytosine with a MIC≥ 2μg/ml, caspofungin with a MIC≥ 16μg/ml, and fluconazole with a MIC≥ 256μg/ml [11]. The phase 2 study suggests that while voriconazole and posaconazole are highly effective, other commonly used antifungal agents, such as amphotericin B and ketoconazole, are less effective [11].

In the current study (2022), it was found that the resistance to itraconazole had increased by 13% compared to the resistance rate in 2006. Additionally, the resistance rate of voriconazole, which was 0% in 2006 and 2011, rose to 9% in 2023. The resistance rates for Posaconazole and isavuconazole, which were both 0% in 2011, increased to 4.7% and 5.8%, respectively, in 2023. Amphotericin B was discontinued in the treatment of falcons against aspergillosis due to its high resistance rate of 80% in 2011.

Itraconazole resistance has been reported in both humans and animals and may result from mutations in the CYP51A gene responsible for the drug target [12]. However, a noticeable change in a decrease to itraconazole resistance was found in 2011 and it was due to the wide use of voriconazole in falcons and thus results in automatically less use of itraconazole (13-15). In-vivo clinical studies have shown that voriconazole is an effective and safe treatment for aspergillosis in falcons [14,15]. Voriconazole has also been found to be the most active drug in many in-vitro studies, with lower MIC values compared to other drugs [16, 17, 18]. However, reports of resistance to voriconazole in some avian isolates of *Aspergillus* species are a growing concern in the management of aspergillosis in birds, including falcons [19, 20]. Voriconazole may not be an appropriate choice for the treatment of Mucor mycosis, as it has been found to be resistant to Mucor species.

Fluconazole is an antifungal medication commonly used in the treatment of human medicine, but its efficacy in treating avian aspergillosis is limited [11,20,21]. There are several reasons why fluconazole may fail in the treatment of falcon aspergillosis. Firstly, *Aspergillus* species in birds are often resistant to fluconazole, and other antifungal medications such as itraconazole and voriconazole are preferred [8]. Additionally, avian aspergillosis is often a systemic disease, meaning that the infection has spread beyond the respiratory tract and into other organs.

Posaconazole is an antifungal medication that belongs to the class of triazole drugs. It is a broad-spectrum antifungal agent that has activity against a wide range of fungal pathogens, including *Aspergillus* species, which can cause aspergillosis in birds, including falcons. Posaconazole has been shown to be effective in the treatment of avian aspergillosis, including in falcons [22]. Recent studies found that a small percentage of the Aspergillus isolates were resistant to posaconazole, with MICs that were higher than the clinical breakpoints for this medication and a study noted that the falcon had been treated with multiple antifungal medications over an extended period, which may have contributed to the development of resistance [23-26].

Isavuconazole is a relatively new antifungal medication that has shown promise in the treatment of aspergillosis in birds, including falcons. Isavuconazole belongs to the class of triazole antifungals, which work by inhibiting the synthesis of ergosterol, a critical component of fungal cell membranes. A few studies have reported the use of isavuconazole for the treatment of aspergillosis in falcons [27,28,29]. While the use of isavuconazole in falcons and other avian species is still relatively limited. MIC studies found that isavuconazole had the lowest MICs among the antifungal drugs tested, indicating that it may be a promising treatment option for aspergillosis in falcons [29]. These results are consistent with previous studies that have shown isavuconazole to be effective against Aspergillus infections in humans and animals. However, this study also identified some isolates with MICs higher than 1 μg/ml, indicating resistance to antifungal drugs, particularly in *A. fumigatus*. Antifungal resistance is a growing problem in both human and veterinary medicine, and the emergence of resistant strains of *Aspergillus* species is a significant challenge to the effective treatment of aspergillosis.

Comparing the MIC values obtained in this study with those reported in the literature for human fungal infections, found that the MIC values for most of the isolates were within the susceptible range for human infections. The median MIC values of voriconazole and itraconazole for *A. fumigatus* isolates in this study were 0.19 μg/ml and 1 μg/ml, respectively, which are lower than the clinical breakpoints for susceptibility in humans, 1 μg/ml, and 2 μg/ml, respectively [30]. However, we observed some differences in MIC values between this study and those reported in the literature for human infections. For instance, the median MIC value of posaconazole for *A. fumigatus* isolates in this study was 0.3 μg/ml, which is slightly higher than the clinical breakpoint for susceptibility in humans, 0.125 μg/ml [30]. Similarly, the median MIC value of isavuconazole for *A. flavus* isolates in this study was 0.38 μg/ml, which is higher than the clinical breakpoint for susceptibility in humans, 0.03 μg/ml [31].

Overall, these results suggest that the antifungal agents tested in this study could be effective in treating fungal infections in falcons caused by the isolates tested. However, caution should be exercised when extrapolating MIC values from veterinary to human medicine, as there can be differences in the pharmacokinetics and pharmacodynamics of antifungal agents between different species. Additionally, it is important to consider other factors such as clinical efficacy and safety when selecting antifungal agents for the treatment of fungal infections in animals. The mic-resistant value to voriconazole is >2 μg/ml in humans but in birds >1 μg/ml is considered as resistant. The efficacy of antifungal drugs may be reduced in cases where *Aspergillus* species are present in the air sac of falcons with high Aspergillus galactomannan antigen levels [31,32]. In conclusion, our study provides valuable information on the susceptibility of various fungal isolates from the air sac of falcons to antifungal agents which are also commonly used in human medicine.

## Conclusion

In conclusion, this study analyzed the minimum inhibitory concentrations (MICs) of antifungal drugs against different fungal isolates from the air sac of falcons. The most isolated fungi were *A. fumigatus, A. flavus, A. niger*, and *A. terreus*, which can cause various infections in both humans and animals, including aspergillosis, which is a major concern in falcons. The study found that isavuconazole, posaconazole, and voriconazole had the lowest MICs among the antifungal drugs tested, suggesting that they could be effective treatment options for aspergillosis in falcons. However, itraconazole, which was widely used to treat aspergillosis in falcons, showed higher MIC values with emerging resistance. The high MICs for some isolates, especially *A. fumigatus*, suggest a risk of antifungal resistance, and thus MIC studies must be followed together with treatment. This study showed a significant increase in resistance to antifungals, with 34% of the isolates being resistant to itraconazole, and resistance rates for posaconazole and isavuconazole rising to 4.7% and 5.8%, respectively, in 2023. Amphotericin B showed 51% resistance in 2006, which rose to 80% in 2011, leading to its discontinuation from treatment and MIC study of falcons.

Overall, this study highlights the significant challenge of emerging antifungal resistance, which is not only a problem in falcon medicine but also in human medicine, as fungal diseases are emerging in association with post-COVID-19. The COVID-19 virus can weaken the immune system, making individuals more susceptible to fungal infections. It is essential to consider these emerging trends and continually monitor the resistance patterns of antifungal drugs to ensure effective treatment of fungal infections in both humans and animals. Additionally, careful consideration of other factors, such as clinical efficacy and safety, is crucial when selecting antifungal agents for the treatment of fungal infections in animals. Lastly, it is important to exercise caution when extrapolating MIC values from veterinary to human medicine, as there may be differences in the pharmacokinetics and pharmacodynamics of antifungal agents between different species.

## Acknowledgments

The authors would like to thank bioMerieux, France, and Al Hayat Pharmaceuticals, UAE for their cooperation in this study.

## Conflict of interest

The authors have no conflicts of interest to declare.

### Funding Source

N/A

### Statement of Ethics

N/A

## Notes

### Competing Interest Statement

The authors have declared no competing interest.

### Summary of Updates

Antifungal resistance is a growing concern in the medical community, as fungal infections are becoming increasingly difficult to treat. In this study, Falco species were used as novel models for studying antifungal resistance since Aspergillosis, a fungal disease is common in falcons. The most isolated fungi in this study were A. fumigatus, A. flavus, A. niger, and A. terreus, all of which can cause aspergillosis in falcons. Isavuconazole, posaconazole, and voriconazole had the lowest MICs among the drugs tested, suggesting that they may be effective treatment options. However, this study showed that 34% of the isolates were resistant to itraconazole, which is an increase from 21% in 2006. There is no resistance to voriconazole found in 2006 and 2011, but a 9% resistance rate was noted in 2022. Similarly, there is no resistance to posaconazole and isavuconazole was noticed in 2011, but the resistance of 4.7% and 5.8%, respectively was noticed in 2022. Amphotericin B, which showed a 51% resistance rate in 2006, became even more resistant with an 80% rate in 2011, leading to its discontinuation from the treatment of falcons against aspergillosis. This study highlights a significant rise in antifungal resistance, which is a challenging problem in both falcon and human medicine.

## References

1. Patrick R. Aspergillosis. Avian Medicine 3^rd^ ed, Samour, 2015, Elsevier, page 460 -471. https://www.elsevier.com/books/avian-medicine/samour/978-0-7234-3832-8

2. Pascal A, Veronica R C, Grégory J, et al. Aspergillosis in Wild Birds, J Fungi (Basel). 2021 Mar; 7(3): 241. doi: 10.3390/jof7030241.

3. Michael PJ, Susan EO. The diagnosis of aspergillosis in birds. Seminars in Avian and Exotic Pet Medicine. 2000, Volume 9 (2), 52–58. https://doi.org/10.1053/AX.2000.4619.

4. Maria EKJ, Susanne V, Julia B. Aspergillosis in Birds: An Overview of Treatment Options and Regimens. Journal of Exotic Pet Medicine. (2015) Volume 24 (3), 296–307. https://doi.org/10.1053/j.jepm.2015.06.012

5. Beernaert LA, Pasmans F, Waeyenberghe LV. Aspergillus infections in birds: a review. Avian Pathology, 2010, Volume 39 (5), https://doi.org/10.1080/03079457.2010.506210

6. Assessment and Clinical implication of Aspergillosis in GYR Falcons (Falco rusticolus). Acta Scientific Veterinary Sciences (ISSN: 2582-3183). 2022. Volume 4 (4), https://actascientific.com/ASVS/pdf/ASVS-04-0356.pdf

7. Amin H, Mohammad HS, Reza D, et al A study on the existence of Aspergillus in birds in the farms around Urmia-Iran. Journal of Stored Products and Postharvest Research Vol. 2(11), pp. 235–236, 2011. https://academicjournals.org/article/article1379949342_Hashempour%20et%20al.pdf

8. Silvanose CD, Bailey TA, Di Somma A. Susceptibility of fungi isolated from the respiratory tract of falcons to amphotericin B, itraconazole and voriconazole: 26 August 2006 https://doi.org/10.1136/vr.159.9.282

9. Grazyna Z, Stanislaw T, Aneta N. Drug resistance of Aspergillus fumigatus strains isolated from flocks of domestic geese in Poland, Poultry Science 2014, 93 (5), 1106–1112, https://doi.org/10.3382/ps.2013-03702

10. Wiederhold NP, Singh N, Gibas C, Lewis RE (2020). Reduced susceptibility to amphotericin B in Aspergillus fumigatus isolates from birds of prey. Veterinary Microbiology, 244, 108679. Doi: 10.1016/j.vetmic.2020.108679

11. Silvanose CD, Bailey TA, Di Somma A, In vitro sensitivity of Aspergillus species isolated from the respiratory tract of falcons - Vet Scan, Online Veterinary Medical Journal, 2020. https://journal.vetscan.co.in/index.php/vs/article/view/106

12. Andréia S, Ana PR, Laura B D, et al, Antifungal susceptibility profile of Aspergillus fumigatus isolates from avian lungs. Pesq. Vet. Bras. 40(2):102-106, February 2020. DOI: 10.1590/1678-5150-PVB-6297. https://www.scielo.br/j/pvb/a/qBjRDnFxR9qRNvTqN7S5XNp/?format=pdf&lang=en

13. Beernaert L A, Pasmans F, Waeyenberghe LV, et al. Avian Aspergillus fumigatus strains resistant to both itraconazole and voriconazole. Antimicrob Agents Chemother. 2009 May;53(5):2199–201. doi: 10.1128/AAC.01492-08.

14. Michael PJ. Selected Infectious Diseases of Birds of Prey. Journal of Exotic Pet Medicine, 2006, Volume 15 (1), 5–17. https://doi.org/10.1053/j.jepm.2005.11.008

15. Antonio Di Somma, Tom Bailey, Christudas Silvanose, et al. The Use of Voriconazole for the Treatment of Aspergillosis in Falcons (Falco Species). J. of Avian Medicine and Surgery, 21(4):307–316 (2007). https://doi.org/10.1647/1082-6742(2007)21[307:TUOVFT]2.0.CO;2

16. Schmidt V, Demiraj F, Di-Somma A, Plasma concentrations of voriconazole in falcons. Vet record.25 August 2007 https://doi.org/10.1136/vr.161.8.265

17. Espinel-Ingroff A, Chowdhary A, Cuenca-Estrella M, et al. Inter-national evaluation of MIC distributions and epidemiological cutoff value (ECV) definitions for Fusarium species identified by molecular methods for the CLSI broth microdilution method. Antimicrob Agents Chemother. 2016;60(2):1079-1084. Doi: 10.1128/AAC.01730-15. PMID: 26643343.

18. Daniel CC, Neil AF. Aspergillosis: update on causes, diagnosis, and treatment. Companion animal. Jan 2016, https://doi.org/10.12968/coan.2016.21.1.50

19. Patrick TR. Comparative Pharmacokinetics of Antifungal Drugs in Domestic Turkeys, Red-Tailed Hawks, Broad-Winged Hawks, and Great-Horned Owls, July 1985, Avian Diseases 29(3):649–61, DOI: 10.2307/1590656.

20. Emerging Fungal Infections: from the Fields to the Clinic, Resistant Aspergillus fumigatus and Dermatophyte Species: a One Health Perspective on an Urgent Public Health Problem, Current Clinical Microbiology Reports (2022) 9:46–51, doi.org/10.1007/s40588-022-00181-3. https://link.springer.com/article/10.1007/s40588-022-00181-3

21. P. Azmanis, L. Pappalardo, Ziad AJS et al, Pharmacokinetics of voriconazole after a single intramuscular injection in large falcons (Falco spp.). Medical mycology 2019. DOI:10.1093/mmy/myz102.

22. Faezezeh M, Seyed J, Seyed MS, et al. Isolation and Characterization of Clinical Triazole Resistance Aspergillus fumigatus in Iran. Iran J Public Health. 2018 Jul; 47(7): 994–1000. PMC6119562.

23. Wellehan JF Jr, Johnson AJ, Zajac AM, et al. In vitro and in vivo antifungal susceptibility of Aspergillus fumigatus isolated from avian species. J Avian Med Surg. 2011;25(4):248-253. Doi: 10.1647/2010-0062.1. PMID: 22330168.

24. Panagiotis A, Lucia P, Ziad AJS, et al, Disposition of posaconazole after single oral administration in large falcons (Falco spp): Effect of meal and dosage and a noncompartmental model to predict effective dosage. Medical Mycology, Volume 59, Issue 9, Sep 2021, 901–908, https://doi.org/10.1093/mmy/myab019.

25. Robert DD, Implication of Mycoses in clinical disorders, Clinical avian Medicine (Ed. Baker JR), 1980. https://avianmedicine.net/wp-content/uploads/2013/08/29_mycosis.pdf

26. Daniel E, Esther S. Diagnostic Aspects of Veterinary and Human Aspergillosis. Front. Microbiol., June 2018. Vol 9 - 2018 | https://doi.org/10.3389/fmicb.2018.01303.

27. Arendrup MC, Jensen RH, Meletiadis J. In vitro activity of isavuconazole against Aspergillus and Candida. J Antimicrob Chemother. 2015;70(10):2918-2923. Doi: 10.1093/jac/dkv197. PMID: 26113596.

28. Jeffrey DJ, Helmut JFS, Juergen P, Spotlight on isavuconazole in the treatment of invasive aspergillosis and mucormycosis: design, development, and place in therapy. Drug Des Devel Ther. 2018; 12: 1033–1044. doi: 10.2147/DDDT.S145545

29. Jesús G, Teresa P, Sandra R, et al. In vitro antifungal activities of isavuconazole (BAL4815), voriconazole, and fluconazole against 1,007 isolates of zygomycete, Candida, Aspergillus, Fusarium, and Scedosporium species. Antimicrob Agents Chemother. 2008 Apr;52(4):1396–400. doi: 10.1128/AAC.01512-07. PMID: 18212101, PMCID: PMC2292541.

30. Arendrup MC, Meletiadis J, Mouton JW, et al. (2017). EUCAST Definitive Document E.DEF Method for the determination of broth dilution minimum inhibitory concentrations of antifungal agents for conidia forming molds. European Committee on Antimicrobial Susceptibility Testing.

31. Nebbia P, Pantchev N, Leisewitz A, et al. (2019). Antifungal drugs in veterinary medicine: an overview. Journal of veterinary pharmacology and therapeutics, 42(2), 129–142. DOI: 10.1111/jvp.12647, PMID: 30714126

32. Williams HL, Wapstra P, Graham J. Aspergillosis in falcons: a review, Veterinary Microbiology: vol. 165, no. 3-4, pp. 181–197, 2013. DOI: 10.1016/j.vetmic.2013.03.002, PMID: 23602181.

